# The periplasmic chaperone Skp prevents misfolding of the secretory lipase A from *Pseudomonas aeruginosa*

**DOI:** 10.1101/2022.03.01.482502

**Authors:** Athanasios Papadopoulos, Max Busch, Jens Reiners, Eymen Hachani, Miriam Bäumers, Lutz Schmitt, Karl-Erich Jaeger, Filip Kovacic, Sander H.J. Smits, Alexej Kedrov

## Abstract

*Pseudomonas aeruginosa* is a wide-spread opportunistic human pathogen and a high-risk factor for immunodeficient people and patients with cystic fibrosis. The extracellular lipase A belongs to the virulence factors of *P. aeruginosa*. The lipase undergoes folding and activation in the periplasm prior the secretion. Here, we demonstrate that the ubiquitous periplasmic chaperone Skp of *P. aeruginosa*, but not SurA, FkpA, PpiD or YfgM, efficiently prevents misfolding of the aggregation-prone lipase A and facilitates its activation by a specific foldase LipH. Small-angle X-ray scattering visualizes the trimeric architecture of *P. aeruginosa* Skp and identifies two primary conformations of the chaperone, a compact and a widely open. We describe two binding modes of Skp to the lipase, with affinities of 20 nM and 2 μM, which correspond to 1:1 and 1:2 stoichiometry of the lipase:Skp complex. Two Skp trimers are required to stabilize the lipase via the apolar interactions, which are not affected by high salt concentrations typical for the sputum of cystic fibrosis patients. The chaperoning effect of Skp points to its potent role in maturation and secretion of the lipase in *Pseudomonas* species.

## Introduction

The Gram-negative bacterium *Pseudomonas aeruginosa* is a wide-spread opportunistic human pathogen of the highest biomedical importance, as indicated by the World Health Organization ^1,2^. The pathogenic potential of *P. aeruginosa* is associated with multiple secreted virulence factors, i.e. the extracellular enzymes, such as exotoxins, lipases and elastases, which facilitate the bacterial infection and adaptation pathways ^3^. The lipase A (LipA; Figure 1A) belongs to the most ubiquitously secreted extracellular enzymes ^4–6^. Secreted LipA is able to hydrolyse long- and short-chain triacylglycerols and, in cooperation with the phospholipase C, it facilitates the release of inflammatory mediators from the host cells ^7^. LipA accumulates in the biofilm matrix of *P. aeruginosa* on infected tissues, where it interacts with the bacterial exopolysaccharide alginate ^8^. Although the physiological role of this interaction is not yet clarified, the biofilm assembly contributes to the bacterial growth, differentiation, and communication within the infection cycle, so the abundant LipA is seen as an element of the bacterial pathogenicity ^8^.

**Figure 1.**
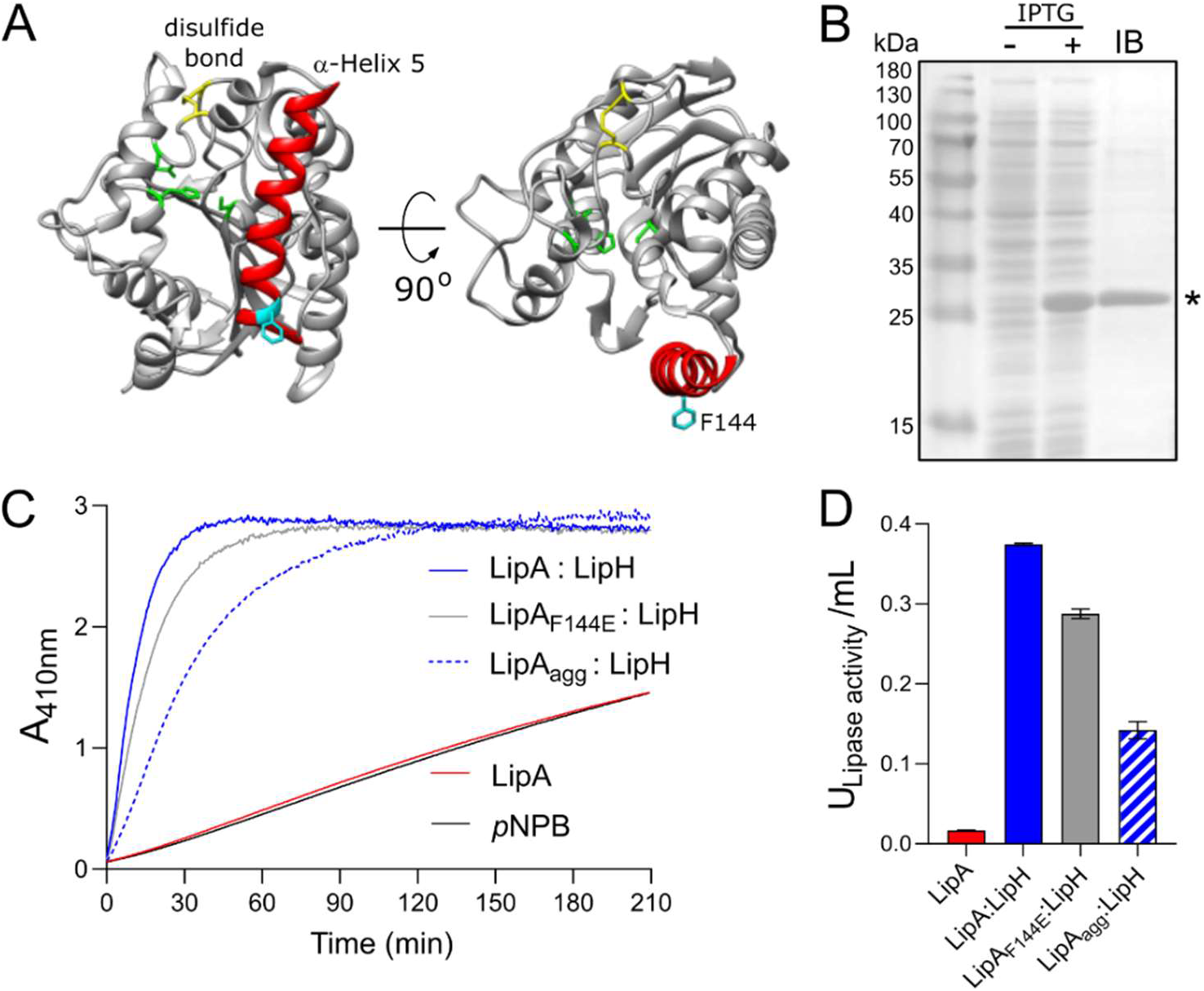
Lipase A from *Pseudomonas aeruginosa* PAO1. **A)** Structure of the lipase LipA (PDB ID: 1EX9). The catalytic triad of Ser-82, Asp-229 and His-251 is shown in green, the disulfide bridge between Cys-183 and Cys-235 is shown in yellow. The α-helix 5 forming the lid domain is shown in red, with the residue F144 in cyan. **B)** LipA expression and purification visualized on SDS-PAGE. Cell lysates prior adding IPTG (“-“) and after 2 hours of expression (“+”) are shown. Lane “IB”: isolated inclusion bodies. LipA band is indicated with asterisk. **C)** Lipase activity *in vitro* mediated by the foldase LipH. Change in absorbance at 410 nm is associated with hydrolysis of the substrate, para-nitrophenyl butyrate (*p*NPB). Once being folded in presence of LipH, the wild-type LipA and the mutant LipA_F144E_ are able to hydrolyse *p*NPB (blue and grey solid traces, respectively). Pre-incubation of the wild-type lipase in the absence of LipH (LipA_agg_) leads to the partial inhibition of hydrolysis (blue dashed trace). LipA activity in the absence of LipH (red trace), and the autohydrolysis of the substrate (black trace) are indicated. **D)** Quantification of the hydrolytic activity of LipA. The lipase activity was calculated based on the colorimetric signal after 15 min of *p*NPB hydrolysis (panel C). The assays were performed in technical triplicates, the mean values and the standard deviations (SD) are shown.

Similar to other secretory proteins, LipA is synthesised as a precursor with an N-terminal signal peptide, a hydrophobic stretch of 26 amino acids, that targets the unfolded lipase to the general SecA:SecYEG translocation pathway ^9^. After the translocation across the cytoplasmic membrane, the signal peptide is cleaved off to release LipA into the periplasm for folding and maturation, followed by export via the type II secretion system (T2SS) ^10^. LipA spontaneously folds into a compact intermediate state that manifests secondary and tertiary structure, however it lacks the enzymatic activity ^11,12^. To achieve the functional conformation prior to T2SS-mediated export, LipA of *P. aeruginosa* requires interaction with a specific foldase LipH encoded in the same operon as LipA ^13,14^. LipH is a membrane-anchored protein with its C-terminal chaperone domain protruding into the periplasm. The available crystal structure of the homologous lipase:foldase complex from *Burkholderia glumae* ^15^ and extensive biochemical and bioinformatics analysis ^11,12,16,17^ suggest that LipH-bound LipA acquires a so-called “open state”, where its α-helix 5 is displaced as a “lid” aside to provide access to the catalytic site. Thus, LipH has been described as a steric chaperone that ensures correct positioning of the LipA lid domain, being a distinct feature of the lipase maturation pathway.

In the absence of the chaperone LipH, the non-activated LipA is prone to aggregation in the crowded bacterial periplasm, which is followed by proteolytic degradation or accumulation in inclusion bodies ^18^. Alike, aggregation and proteolysis challenge the biogenesis of ubiquitous outer membrane proteins (OMPs), which, once translocated from the cytoplasm, should cross the periplasm prior to the integration into the outer membrane ^19,20^. To facilitate their targeting and disaggregation, a range of ATP-independent chaperones is present in the periplasm at micromolar concentrations ^9,21^. Widely conserved proteins SurA, Skp, and FkpA are the best-studied chaperones, which serve as holdases and escort client proteins to the outer membrane, while preventing their premature folding or aggregation. The soluble chaperones are further complemented by PpiD and YfgM proteins anchored at the inner membrane and associated with the SecYEG translocon ^22,23^. Notably, the periplasmic chaperones are also involved in biogenesis of several secretory proteins, including a bacterial lipase ^24–27^, but the mechanism of these interactions has been barely investigated.

Here, we demonstrate that folding of LipA is challenged by the high aggregation propensity of its lid domain, while a mutation within this region as well as interactions with the foldase LipH greatly stabilize the protein. When examining interactions of LipA with the general periplasmic chaperones of *P. aeruginosa*, we discover a potent anti-aggregation effect of the chaperone Skp, while LipA:Skp interactions do not prevent LipH-mediated activation of the lipase. The structural analysis of Skp reveals a characteristic jellyfish-shaped trimeric assembly and suggests extensive conformational dynamics. The hydrophobic central cavity within the Skp trimer may expand largely to accommodate a LipA molecule, although two bound copies of Skp trimer are required to prevent LipA from aggregation. The results suggest that Skp may be a potent chaperone in the LipA biogenesis pathway that ensures client’s maturation under challenging environmental conditions.

## Results

### LipA aggregation is mediated by the lid domain

To investigate folding and activation of the lipase *in vitro*, LipA lacking the N-terminal signal peptide was heterologously overexpressed in *E. coli*. The overexpression resulted in formation of inclusion bodies consisting nearly exclusively of LipA, and the protein was isolated in the urea-denatured state (Figure 1B). The denatured LipA is a relevant mimetic of the protein that enters the periplasm as an unfolded polypeptide chain, and it was used for the further analysis. To examine whether the recombinant protein may be refolded into its functional form, LipA was diluted into the urea-free low-salt TGCG buffer (5 mM Tris pH 9.0, 5 mM glycine, 1 mM CaCl_2_ and 5% glycerol) ^6^ and incubated in the presence of the foldase chaperone LipH. The enzymatic activity of LipA was assessed then via measuring the hydrolysis of a model substrate, para-nitrophenyl-butyrate (*p*NPB): Accumulation of the product, p-nitrophenolate, was followed colorimetrically until reaching the signal saturation, and the activity of the lipase A was quantified (Figures 1C and D). In the absence of LipH, the signal remained at the level of the autohydrolysis of *p*NPB, so the hydrolytic activity of LipA was not induced. Thus, the recombinant LipA could be folded *in vitro*, and LipH was required for activation of LipA.

Notably, if LipA was incubated in the urea-free buffer prior to adding the foldase LipH, its hydrolytic activity was substantially reduced, as the amount of p-nitrophenolate generated in the first 15 min of the reaction decreased three-fold (Figure 1C and D). We speculated that the lipase underwent aggregation/misfolding in the absence of the foldase LipH, causing the loss of activity. Indeed, approx. 50 % of LipA could be sedimented upon mild centrifugation at 21000 g (Figure 2A). The sedimentation was enhanced upon increasing the ionic strength of the solution, reaching 70 % and 95 % in the presence of 50 and 150 mM NaCl, respectively (Figure 2B) and abundant large particles were observed by negative-stain electron microscopy (Suppl. Figure 1A). The stoichiometric amount of LipH stabilised the lipase, although the efficiency decreased at the elevated salt concentrations (Figure 2B). Since LipA:LipH binding is primarily driven by electrostatic interactions ^11,12,15^, the affinity of the complex was likely reduced at the elevated ionic strength and LipA aggregation became a dominant pathway even in the presence of the chaperone.

**Figure 2.**
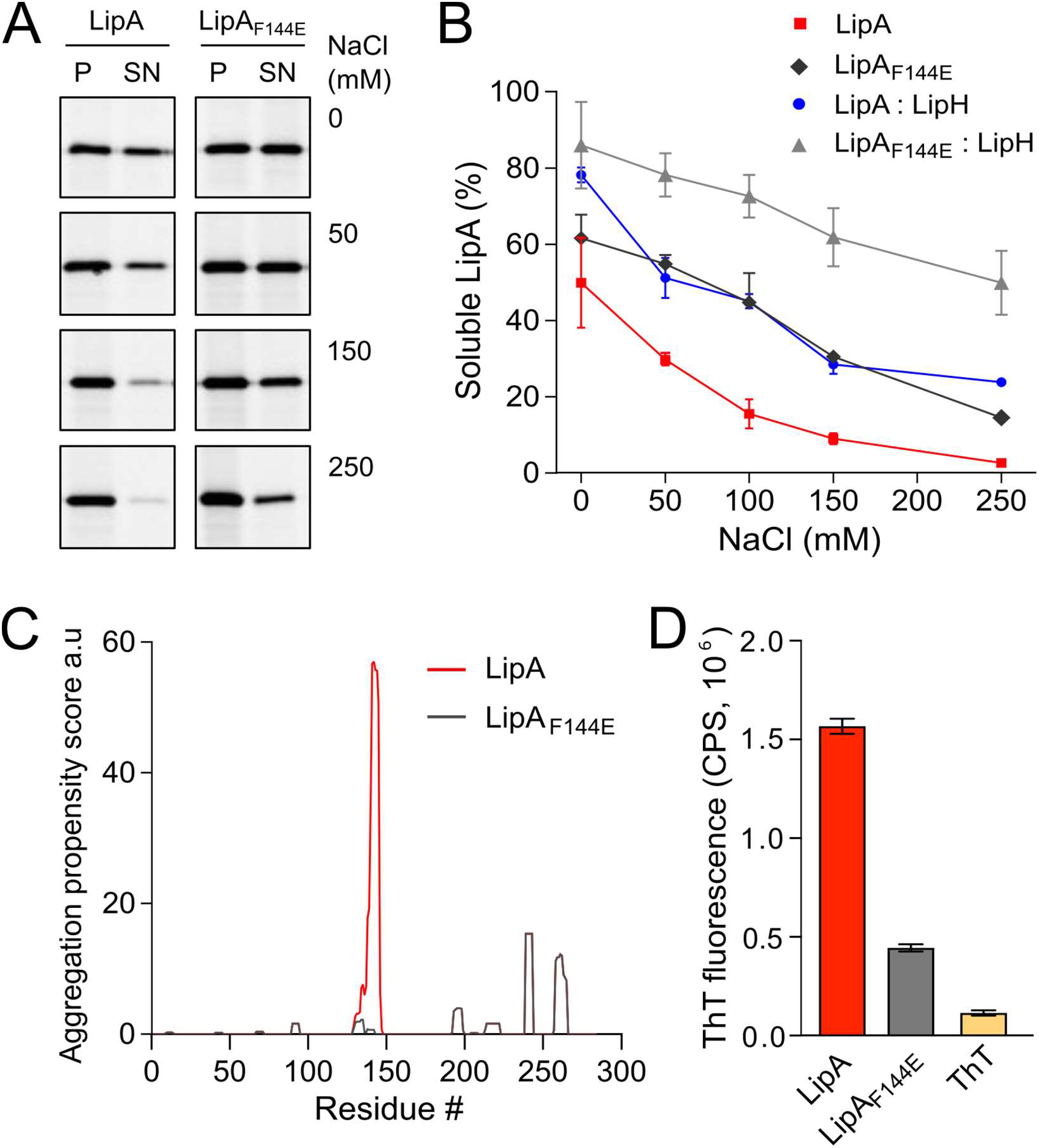
LipA aggregation propensity is sensitive to the environment and the structure of the lid domain. **A)** SDS-PAGE of the aggregated and soluble LipA variants separated into the pellet (P) and the supernatant (SN) fractions, respectively, upon centrifugation. The concentrations of NaCl in the buffer are indicated. For the in-gel fluorescence detection, the lipases were labelled with fluorescein-5-maleimide. **B)** Quantification of the soluble LipA at various conditions. The fraction of the soluble wild-type LipA and the mutant LipA_F144E_ at various salt concentrations and, optionally, in presence of LipH was calculated from SDS-PAGE (panel A). The assays were performed in technical triplicates, the mean values and SD are shown. **C)** Sequence-based prediction of LipA aggregation propensity by TANGO algorithm ^28^. Profiles for the β-aggregation propensity scores of the wild-type LipA (red) and the mutant LipA_F144E_ (grey) indicate the aggregation-prone regions within the polypeptide chain. The aggregation propensity of the wild-type LipA is dominated by the α-helix 5 that forms the lid domain. **D)** LipA aggregation is associated with the formation of β-structured amyloid-like aggregates. The intensity of the amyloid-sensitive ThT fluorescence at 485 nm was measured in the presence of either the wild-type LipA (red), the mutant LipA_F144E_ (grey), or the dye alone (yellow). The assays were performed in technical triplicates, the mean values and SD are shown.

Based on its amino acid sequence, the α-helix 5 known as the “lid domain” of LipA is predicted to be the primary aggregation-prone region within the protein, with a high propensity to misfold into β-strands (Figure 2C, red line) ^28^. Previous studies on a homologous lipase demonstrated that a point mutation Phe to Glu within the lid domain diminished the aggregation propensity and rendered an aggregation-resistant protein ^29^. Based on the structure homology, we recognised an identically positioned phenylalanine residue within *P. aeruginosa* PAO1 LipA at position 144, where the point mutation to glutamate could suppress the lipase misfolding (Figure 2C, black line). To examine the effect of the mutation on the lipase folding and activity, the mutant LipA_F144E_ was overexpressed and isolated in its urea-denatured state. In the presence of LipH, LipA_F144E_ displayed the enzymatic activity, indicating that the mutation did not hinder the chaperone-mediated folding (Figure 1C and D). Indeed, the lid domain is not involved in LipA:LipH contacts ^15^, and the mutated residue is oriented toward the solvent and does not belong to the moiety of the catalytic site (Figure 1A).

To examine whether LipA_F144E_ is indeed more resistant against aggregation than the wild-type lipase, its solubility was probed over a range of salt concentrations. The point mutation had a notable effect on the protein stability: Nearly 50 % of LipA_F144E_ remained soluble in the presence of 100 mM NaCl, where the soluble fraction of the wild-type LipA did not exceed 15 % (Figure 2B). As the aggregated lipase tends to form amyloid-type structures ^29^, we further examined this propensity for the wild-type LipA and LipA_F144E_ mutant. To do so, we measured the fluorescence intensity of the dye thioflavin T (ThT), which fluorescence increases manifold upon binding to β-strands within amyloid-type aggregates ^30^. Once the dye was added to the wild-type lipase in the TGCG buffer supplemented with 20 mM NaCl, its fluorescence intensity increased 10-fold, suggesting that β-stranded structures were formed even at the low salt concentration (Figure 2D and Suppl. Figure 2B). In contrast, the ThT fluorescence was only modestly affected in the presence of LipA_F144E_, increasing 3-fold above the signal of free ThT, in agreement with the reduced aggregation of the mutant.

As both the mutation within the lid domain of LipA and the LipA:LipH assembly favoured the soluble state of the lipase, we questioned whether both cases involve the same stabilisation mechanism. To address this, the solubility of the wild-type LipA and LipA_F144E_ in the presence of the foldase was compared (Figure 2B). Strikingly, the solubility of the mutant LipA_F144E_ was further enhanced by LipH: At elevated salt concentrations, the fraction of the soluble LipA_F144E_ exceeded approx. two-fold that of the wild-type LipA, so even at 250 mM NaCl the major fraction of LipA_F144E_:LipH remained soluble. Conclusively, the effects of the mutation within the lid domain and the lipase:foldase binding were additive, and distinct mechanisms contributed to the lipase stabilisation.

### Characterisation of periplasmic chaperones of P. aeruginosa

The extensive aggregation of the wild-type LipA observed already at moderate salt concentrations is a potential challenge for its timely interactions with the stabilising foldase LipH, especially under high-salinity conditions within cystic fibrosis patients’ sputum ^31^. As the highly abundant periplasmic chaperones may facilitate folding and secretion of several recombinant proteins, including a lipase of *Burkholderia* ^24–27,32^, we speculated that the chaperones of *P. aeruginosa* may recognize LipA and protect it from aggregation prior interactions with the specific foldase LipH.

To characterize LipA:chaperone interactions in a well-defined *in vitro* system, the primary soluble chaperones Skp, FkpA and SurA lacking their signal peptides (Figure 3A), as well as the periplasmic domains of YfgM and PpiD of *P. aeruginosa* PAO1 were heterologously expressed and purified from *E. coli*. The apparent molecular weights of the chaperones observed in SDS-PAGE agreed with the values calculated for individual protomers (18 kDa for Skp, 20 kDa for YfgM, 27 kDa for FkpA, 47 kDa for SurA and 67 kDa for PpiD; Figure 3B). To scrutinise the oligomeric states of the isolated chaperones, their native masses were determined by size exclusion chromatography combined with multi-angle light scattering analysis (SEC-MALS, Figure 3C). The experiment revealed monomers of SurA (determined molecular weight of 49.6±0.6 kDa), dimers of FkpA (53.2±0.9 kDa; FkpA_2_), and trimers of Skp (58.6±0.7 kDa; Skp_3_), in agreement with the known structures of homologs from *E. coli* ^33–35^. The periplasmic domains of both YfgM and PpiD were found to be monomers, with molecular masses of 22.2±0.4 kDa and 72.2±0.2 kDa, respectively. The periplasmic chaperone domain of LipH appears to be monomeric in solution (determined molecular weight 39.4±0.3 kDa). The attempts to analyse the molecular mass of either wild-type LipA or LipA_F144E_ were not successful, as the hydrophobic protein was possibly interacting with the column matrix and could not be eluted with aqueous buffers.

**Figure 3.**
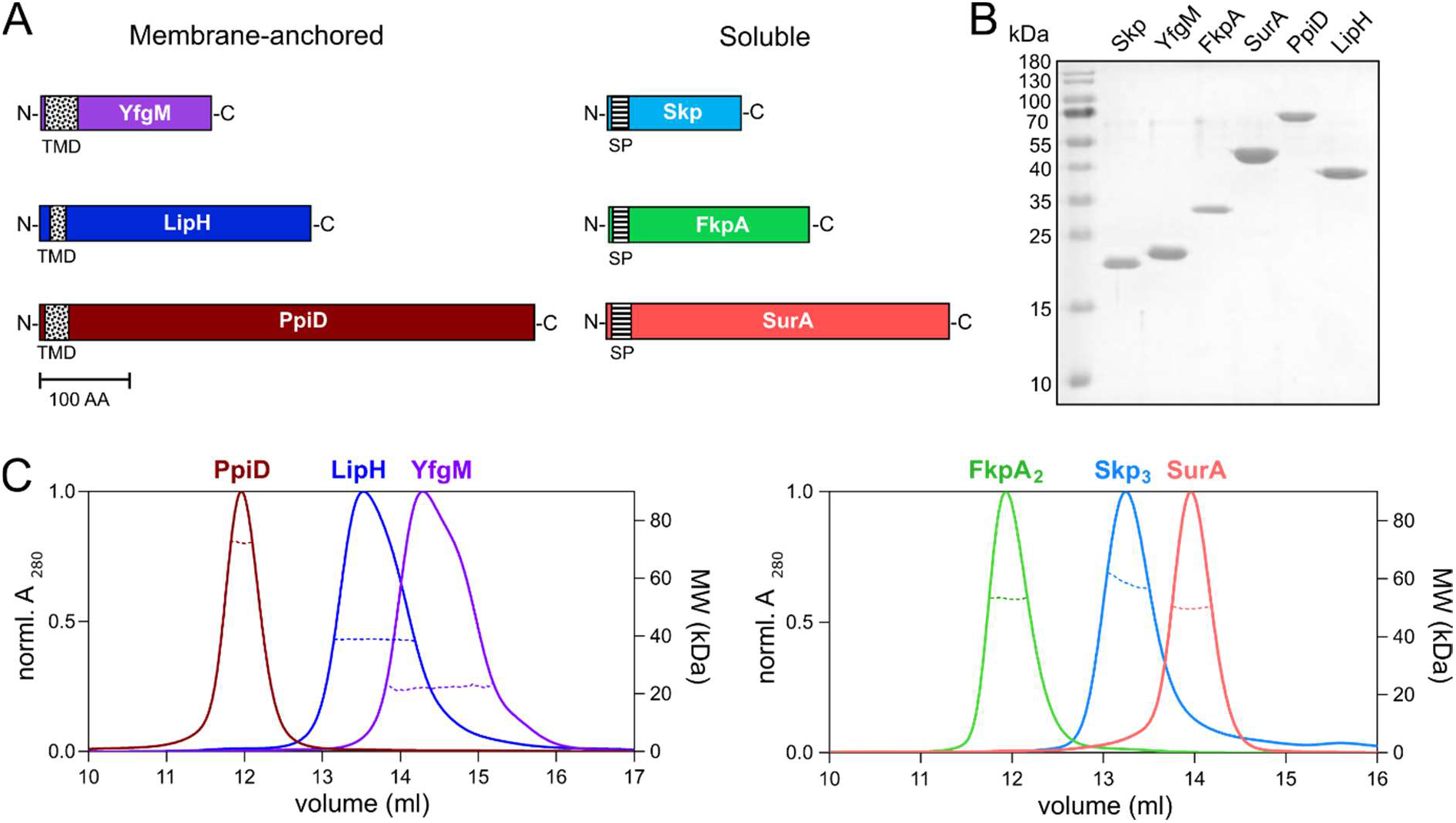
Periplasmic chaperones of *P. aeruginosa* PAO1. **A)** Schematic overview of the periplasmic chaperones of *P. aeruginosa* PAO1. The predicted transmembrane domains (TMD, black-dotted) and the signal peptides (SP, black-ruled) were removed to express the soluble chaperones. Scale bar: 100 amino acid residues (AA). **B)** SDS-PAGE of the isolated periplasmic chaperones of *P. aeruginosa*. **C)** SEC-MALS analysis of the periplasmic chaperones to determine the oligomeric states in solution. Solid traces represent the absorbance at 280 nm, dashed traces indicate the molecular weights (y-axis on the right) calculated from the light scattering. The analysis reveals dimers of FkpA (FkpA_2_) and trimers of Skp (Skp_3_), while SurA, YfgM, PpiD and LipH are monomeric in solution.

### The chaperone Skp prevents LipA from aggregation

The propensity of the individual chaperones to prevent LipA aggregation was first evaluated by the sedimentation analysis. To mimic the natural abundance of SurA, FkpA_2_ and Skp_3_ in the bacterial periplasm, 5 μM of chaperones were used in the experiments ^9,21^, corresponding to the lipase:chaperone molar ratio 1:10, and the salt concentration was set to 150 mM NaCl. In the absence of the chaperones, only 10 % of LipA was found in the soluble form, and the value increased moderately in the presence of either SurA or FkpA_2_ (solubility of 15 % and 29 %, respectively; Figure 4A and B). In contrast, Skp_3_ demonstrated prominent stabilisation of LipA, as the soluble fraction reached 66 %. In the presence of the lipase-specific foldase LipH, more than 80 % LipA was found soluble (LipA:LipH molar ratio 1:1).

**Figure 4.**
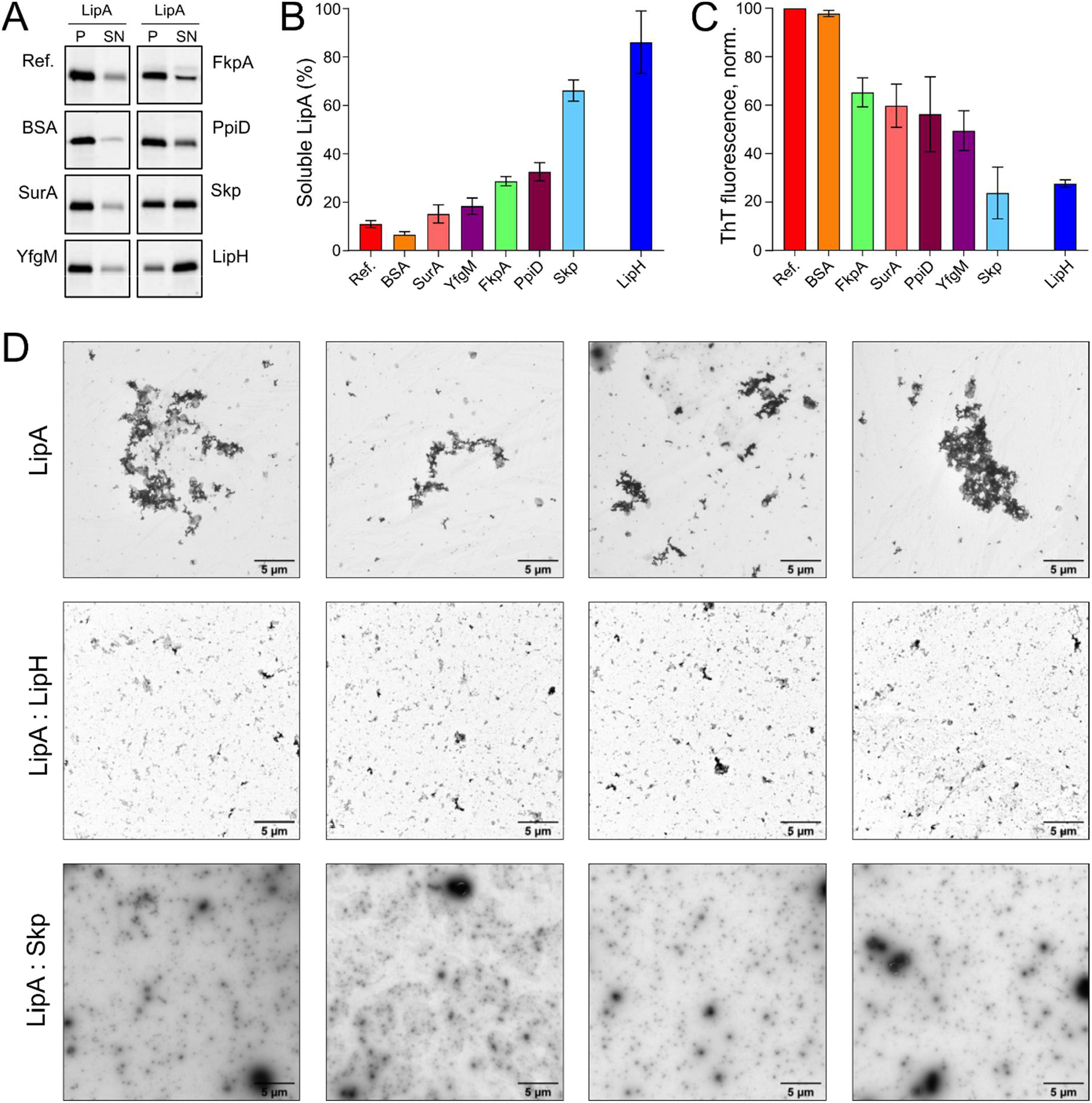
The periplasmic chaperone Skp rescues LipA from aggregation. **A)** LipA solubility in the presence of the periplasmic chaperones. The aggregated and soluble LipA fractions are separated in pellet (P) and supernatant (SN) fractions in the sedimentation assay and visualized on SDS-PAGE via in-gel fluorescence. The supplemented chaperones are indicated. Bovine serum albumin (BSA) was used as a negative control. LipA in absence of chaperones is indicated as “Ref.”. **B)** Quantification of the soluble LipA in the presence of the periplasmic chaperones from SDS-PAGE (panel A). The assays were performed in technical triplicates, the mean values and SD are shown. **C)** The periplasmic chaperones suppress formation of LipA β-structured aggregates. ThT fluorescence was recorded for the wild-type LipA in the presence of the periplasmic chaperones and normalized to the fluorescence of LipA alone. The assays were performed in technical triplicates, the mean values and SD are shown. **D**: LipA aggregation in presence and absence of LipH (LipA:LipH) or Skp (LipA:Skp) visualized via negative-stain TEM. Scale bars are indicated.

The membrane-anchored proteins PpiD and YfgM are the recently described chaperones associated with the SecYEG translocon of *E. coli* ^23,36^. Being proximal to the translocon, the peptidyl prolyl isomerase PpiD and the tetratricopeptide repeat-containing protein YfgM of *P. aeruginosa* may interact with LipA at the early stage of its translocation into the periplasm and mediate the handover to the membrane-anchored LipH. Similar to the soluble chaperones above, the potential effects of the periplasmic domains of PpiD and YfgM on LipA aggregation were investigated. YfgM did not stabilize LipA, as the soluble fraction remained below 20 % (Figure 4A and B). This lack of the holdase activity is in agreement with earlier experiments, where luciferase and the precursor of OmpC were tested as putative YfgM clients ^23^. In the presence of PpiD, the soluble fraction of LipA increased up to 34 %, which matched closely the value measured for another peptidyl prolyl isomerase, FkpA. Overall, the sedimentation assay identified the non-specific soluble chaperone Skp as a potent stabilising factor of the lipase.

Next, we questioned whether the chaperones may be capable of preventing the assembly of amyloid-type aggregates of LipA. To probe this putative effect, ThT fluorescence was measured for LipA in the presence of individual chaperones and corrected to the values of the chaperones alone (Suppl. Figure 2). Although most of the chaperones had moderate to low effect on LipA solubility in the sedimentation assay (Figure 4B), they all could reduce the ThT fluorescence at least by 40 %, as compared to LipA alone or LipA in the presence of bovine serum albumin (BSA) (Figure 4C). These values reflect the partial ability of chaperones to prevent amyloid-like structure formation by LipA. Notably, the most prominent effect was observed again for Skp, as the amyloid-specific ThT signal in the presence of the chaperone was reduced by approx. 75 %. Similar low ThT fluorescence intensity was recorded in the presence of LipH.

Finally, negative-stain electron microscopy was used to visualize LipA aggregation. At the elevated ionic strength and in the absence of urea, LipA readily formed micrometer-sized aggregates, commonly assembled in larger clusters (Suppl. Figure 1A and Figure 4D). The sample morphology was altered when either LipH or Skp_3_ were added at the molar ratios of 1:1 and 1:10, respectively, so only sub-micrometer featureless particles could be observed, suggesting reduced aggregation propensity. Thus, we concluded that both Skp and LipH offer protection from the assembly of aggregates, despite different mechanisms of interaction with the lipase.

### The periplasmic chaperones modulate the activation of LipA

The interactions of the periplasmic chaperones with LipA and their anti-aggregation effects may affect the functionality of the lipase, e.g. by directly altering its folding pathways and/or by competing with the specific foldase LipH. None of the general periplasmic chaperones alone rendered the active lipase, as no enzymatic hydrolysis of *p*NPB was observed in the absence of LipH (Figure 5A). To examine the potential competition between the chaperones, we focused on the LipH-mediated activation of LipA in the presence of the general periplasmic chaperones, as it occurs in living cells. The wild-type LipA was pre-incubated in the presence or absence of individual chaperones under conditions of moderate ionic strength (50 mM NaCl, 1 mM CaCl_2_ and 20 mM Tris-HCl pH 8.0) sufficient to induce the lipase aggregation. After the pre-incubation phase, the foldase LipH was added to activate LipA, and the resulting hydrolytic activity against *p*NPB was determined (Figure 5B). The conditions of the pre-incubation phase had clear influence on the enzymatic activity of LipA: In the sample to which no chaperones were initially added, LipA activity was suppressed to a large extent (blue bars in Figures 5A vs 5B), in agreement with the documented aggregation of the lipase. Notably, the pre-incubation in the presence of SurA, FkpA_2_, YfgM or PpiD chaperones led to further two- to four-fold decrease in LipA activity (Figure 5B). Differently though, pre-incubation of LipA with Skp_3_ resulted in modest stimulation of the substrate hydrolysis, in agreement with the antiaggregation effect of the chaperone.

**Figure 5.**
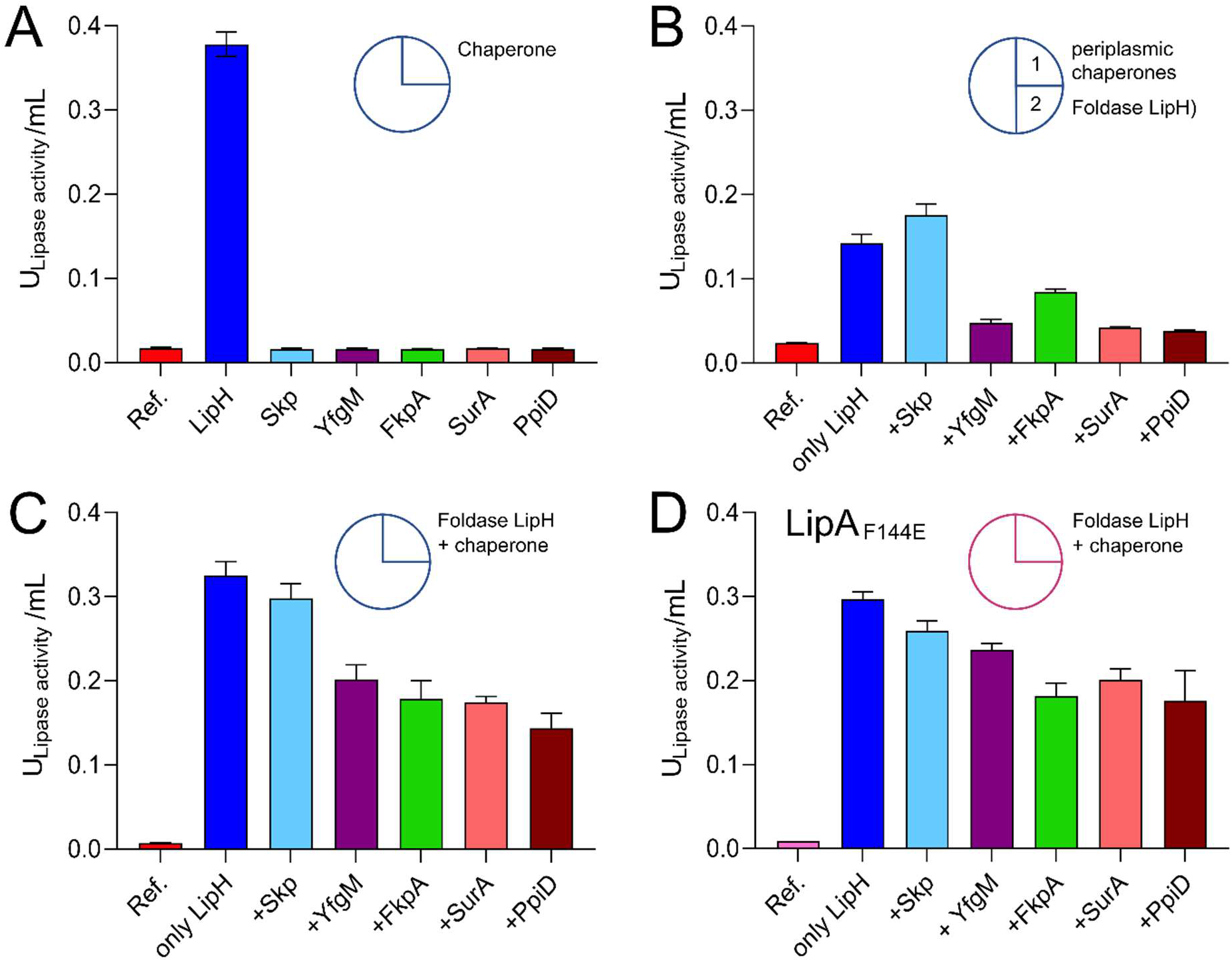
Interplay of periplasmic chaperones upon lipase activation. The lipase activity against *p*NPB over the first 15 min of the hydrolysis reaction is shown in bars. Circle scheme describes the incubation steps. LipA activity in absence of chaperones is indicated as “Ref.”. **A)** Incubation of the wild-type LipA with either the foldase LipH or general periplasmic chaperones. **B)** Initial incubation of the wild-type LipA with general periplasmic chaperones (1) followed by incubation with LipH (2). **C)** Simultaneous incubation of the wild-type LipA with LipH and periplasmic chaperones. **D)** Simultaneous incubation of the mutant LipA_F144E_ with LipH and periplasmic chaperones. The assays were performed in technical triplicates, the mean values and SD are shown.

The reduced LipA activity in the presence of FkpA, SurA, YfgM or PpiD correlated with their poor propensity to prevent LipA aggregation, but also suggested the interference with the LipH-mediated activation, potentially by competing for the lipase binding. To examine this scenario, both, the general chaperones and LipH, were added to LipA simultaneously at the LipA:LipH:chaperone molar ratio of 1:1:10, so the lipase activity was dependent on the folding/aggregation balance and the competition between the chaperones for the lipase. In the presence of either SurA, FkpA_2_, YfgM or PpiD, the activity of LipA dropped by 30-50% (Figure 5C), while only minor, below 10 % inhibition of substrate hydrolysis was observed for LipA:LipH:Skp_3_ sample. Finally, to scrutinise the effect of the chaperone competition, the assay was conducted using the aggregation-resistant LipA_F144E_ mutant (Figure 5D). When either SurA, FkpA_2_, YfgM or PpiD were present in addition to LipH, the lipase activity reduced by 20 to 30 %, so we concluded that the chaperones interfered with the LipH:LipA assembly. The effect of Skp_3_ on LipH-mediated activation was minimal, so the chaperone allowed efficient LipH:LipA interactions, while being capable of preventing LipA aggregation.

### Trimeric Skp of P. aeruginosa acquires multiple conformations in solution

As the biochemical experiments highlighted a potent role of Skp in LipA folding, we set out to shed more light on the functional mechanism of the chaperone. The well-studied Skp homolog from *E. coli* (*Ec*Skp) forms a “jellyfish”-like trimer that resembles the arrangement of prefoldin-type chaperones of mitochondria and archaea ^34,37^. Each protomer consists of the distinct “head” and “arm” domains. The “head” domain of *Ec*Skp is predominantly formed by β-sheets which drive the oligomerization, and the “arm” domain is a helical section forming a hairpin extension. The client-binding pocket between the “arm” domains is constituted by all three promoters. The crystal structures and small-angle X-ray scattering (SAXS) experiments predicted a broad range of conformations for the *Ec*Skp trimer in its apo-state in solution, where three major states termed as “closed”, “intermediate” and “open” were described ^38^.

Mature Skp of *P. aeruginosa* shares 22 % sequence identity with *Ec*Skp and contains three unique proline residues per protomer, including two prolines in the predicted “head” domain and one in each helical “arm”, which may alter the flexibility of the chaperone (Suppl. Figure 3). Thus, we employed SAXS to experimentally assess the conformational ensemble of *P. aeruginosa* Skp in solution (Figure 6A and Suppl. Table 1). In agreement with SEC-MALS data (Figure 3), Skp was found to be a trimer (Skp_3_). The characteristic “jellyfish”-like shape with the “head” and “arm” domains of the trimer could be recognized in the *ab initio* reconstitution of the oligomer structure using the GASBOR program ^39^ (χ^2^ =1.016) (Figure 6A and B). A structural model of P. aeruginosa Skp_3_ built by AlphaFold 2 algorithm ^40,41^ could be fitted into the experimental SAXS envelope (Figure 6B). However, the measured radius of gyration (*R*_g_) of 3.50 nm for the averaged SAXS data is slightly higher than *R_g_* of 3.32 nm calculated via CRYSOL for the modelled structure (χ^2^ =1.28) ^42^, so Skp_3_ of *P. aeruginosa* adopts a more open conformation.

**Figure 6.**
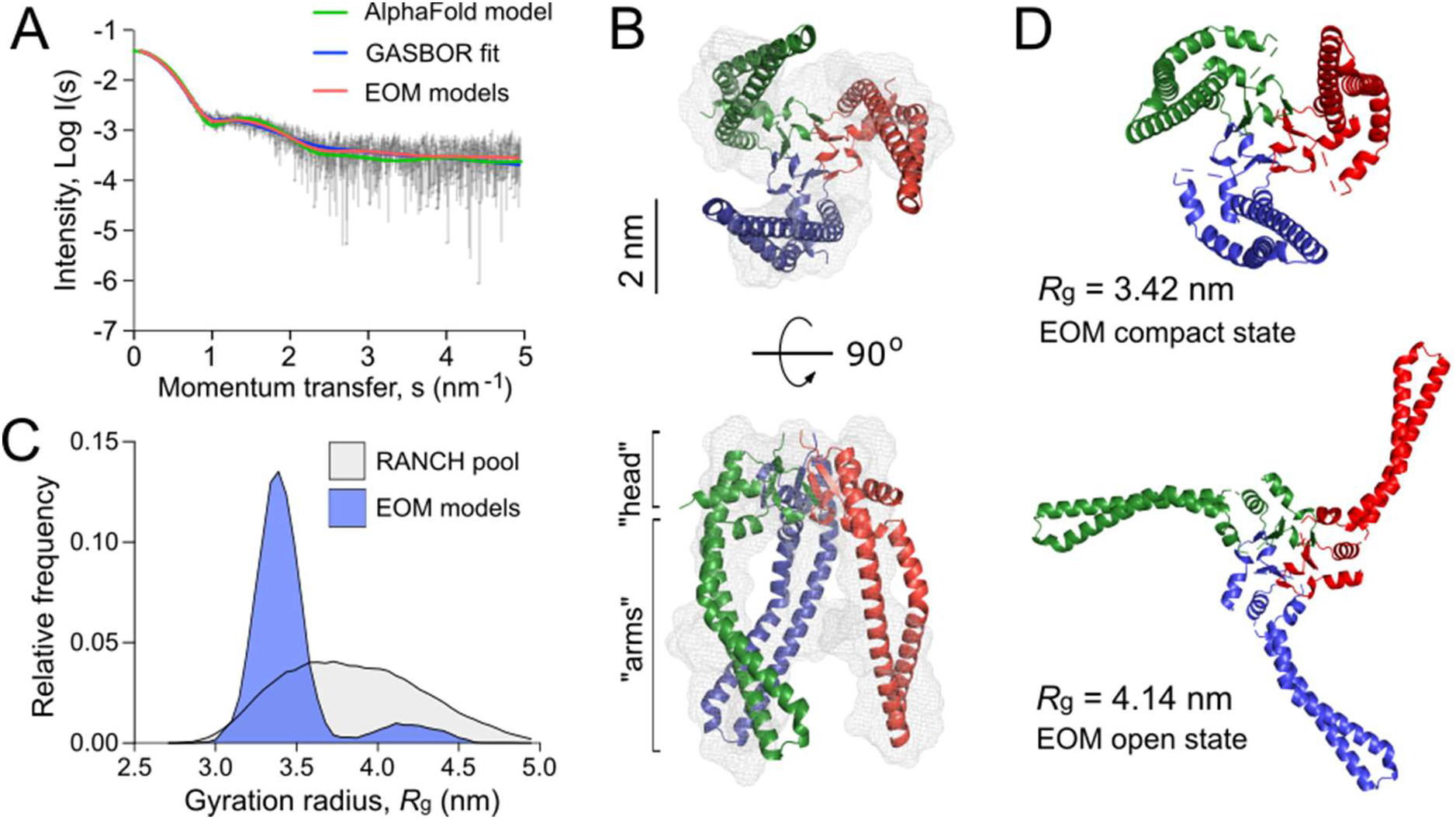
*P. aeruginosa* Skp forms a flexible trimer in solution. **A)** Experimental SAXS data curve is shown in grey dots with error bars. The intensity (log I) is displayed as a function of the momentum transfer (s). The intensity patterns of Skp_3_ models derived from GASBOR (blue), AlphaFold (green), and EOM (red) algorithms are overlaid. **B)** The experimental SAXS envelope of *P. aeruginosa* Skp (grey mesh) calculated using GASBOR algorithm reflects the trimeric structure of the chaperone. The structural model of Skp_3_ is superimposed with the SAXS envelope, with each protomer coloured individually (red, green, blue). The “head” and “arm” structural domains of Skp are indicated. The scale bar is shown on side. **C)** The distribution of Skp_3_ conformations differing by their gyration radii (*R*_g_) calculated by EOM (blue). The original RANCH pool based on 10000 random models is shown in grey. **D)** Representative conformations of *P. aeruginosa* Skp_3_ calculated by EOM in the compact (*R*_g_ of 3.42 nm, *D_max_* of 9.73 nm; fraction ~0.62) and the open (*R*_g_ of 4.14 nm, *D_max_* of 13.63 nm; fraction ~0.12) states.

The SAXS data also revealed some flexibility in Skp_3_ of *P. aeruginosa*, like previously observed for the *Ec*Skp homolog ^38^. To characterize this conformational space, Ensemble Optimization Method (EOM) was applied ^43,44^. EOM generates a pool of sequence- and structure-based models (Figure 6C, RANCH models), and the implemented genetic algorithm GAJOE (Genetic Algorithm Judging Optimisation of Ensembles) selects an ensemble until an optimal fit to the experimental SAXS curve is reached. The selected models describe best the experimental SAXS data, and likely represent the most frequent conformations of the studied protein (Figure 6B). To model the Skp_3_ dynamics by EOM, the “head” domain within the trimer was fixed and the helical “arms” were allowed moving to fit the SAXS data. As a result, the selected ensemble matched the experimental SAXS curves with χ^2^ =0.954 (Figure 6A, EOM models). The distribution of the models showed two primary states, resembling those reported for *Ec*Skp_3_ ^38^: The “compact” state with *R*_g_ of 3.42 nm and maximal dimensions (*D*_max_) of 9.73 nm, which accounts up to 62% of the population, and the “open” state (*R*_g_ = 4.14 nm, *D*_max_ = 13.63 nm; 12% occupancy) (Figure 6D). While the dimensions of the open state were nearly identical for both Skp_3_ homologs, the compact state of Skp_3_ of *P. aeruginosa* was wider than the “intermediate” conformation of *Ec*Skp_3_ (*R*_g_ of 3.42 nm vs. 3.2 nm), and no “closed” state was observed for Skp_3_ of *P. aeruginosa*. Nevertheless, the ability of Skp_3_ of both *E. coli* and *P. aeruginosa* to undergo a large conformational change from a compact to a widely open state suggests that the proteins likely follow the same functional mechanism.

### Two copies of Skp are required to stabilize soluble LipA via apolar contacts

The documented clients of Skp in the bacterial periplasm span a broad range of molecular masses, from 19 to 89 kDa, and they belong to various classes, both membrane-associated and globular proteins ^45,46^. Notably, the stimulatory effect of Skp on secretion of a bacterial lipase has been previously demonstrated, but it remained obscure whether the stimulation was indeed a result of direct client:chaperone interactions ^27^. Our biochemical data substantiates the view that Skp_3_ is a potent chaperone that prevents the lipase aggregation, and here we set out to characterize LipA:Skp_3_ interactions in detail.

Firstly, we questioned whether the interaction of *P. aeruginosa* Skp_3_ with LipA is indeed of hydrophobic nature, as each of the “arms” within the Skp trimer has an apolar side (Figure 7A) that may bind unfolded and/or misfolded clients. As shown for the LipA:LipH complex, elevated ionic strength of the aqueous solution severely destabilizes the hydrophilic binding interface, so the released LipA irreversibly aggregates (Figure 2B). Hydrophobic interactions, in contrast, should be insensitive to the ionic strength and may facilitate the chaperoning at high salt concentrations. The solubility of LipA in the presence of Skp_3_ at the molar ratio 1:10 was examined over a range of NaCl concentrations via the sedimentation assay (Figure 7B). Different to LipA alone and LipA in the presence of LipH, no decay in the soluble fraction was observed when Skp_3_ was present. At the highest examined concentration of 250 mM NaCl, Skp_3_ ensured the lipase solubility of approx. 70 %, while the chaperone-free LipA was nearly completely aggregated, and only 30 % of LipA was found soluble in the presence of LipH. This observation implied that the lipase is recognized via non-polar contacts, likely associated with the unfolded state.

**Figure 7.**
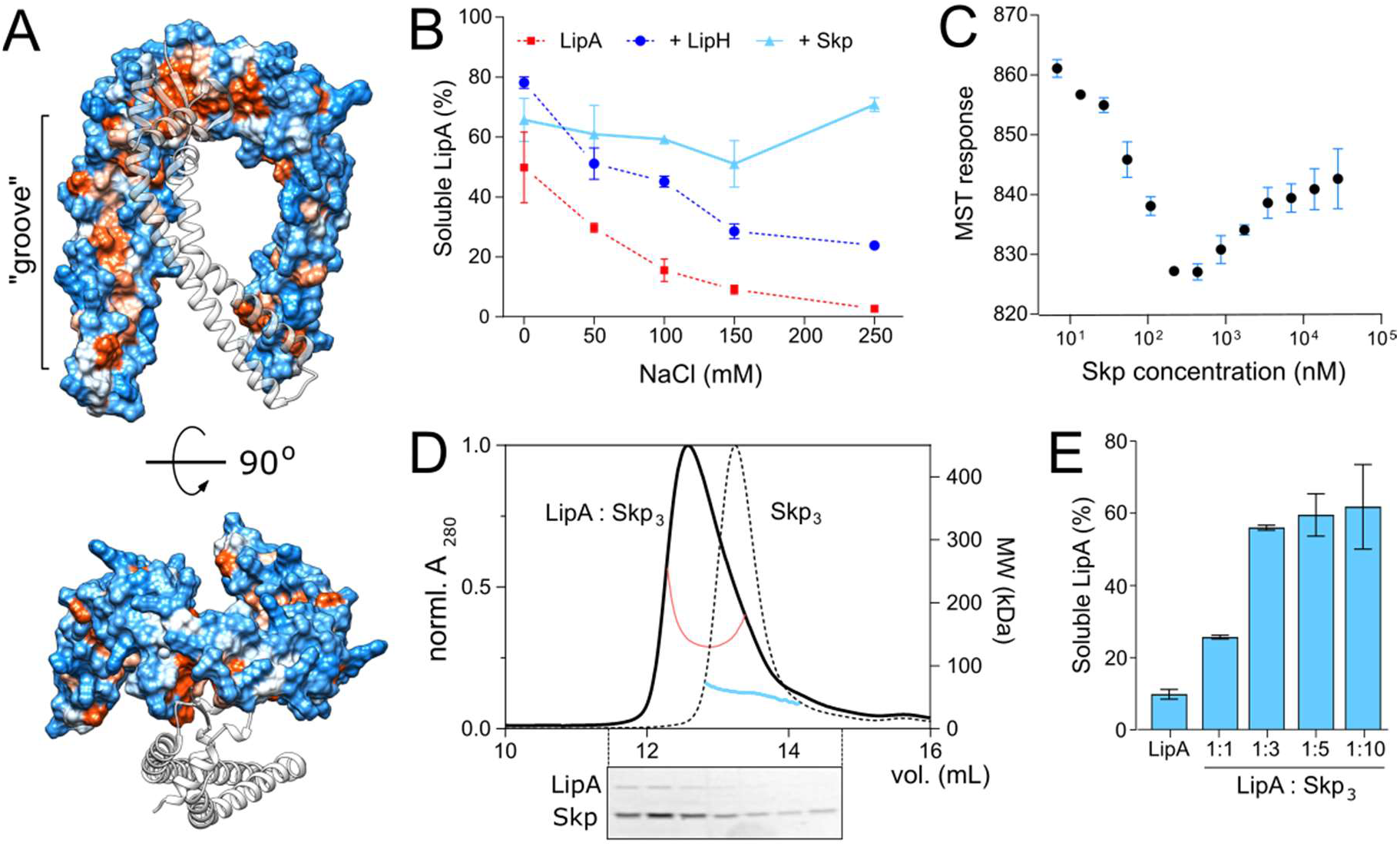
Characterisation of LipA:Skp_3_ interactions. **A)** Surface hydrophobicity plot of *P. aeruginosa* Skp_3_. The hydrophobic groove along the “arm” domain is indicated. Polar/charged residues are shown in blue, apolar in orange. One subunit within the Skp trimer is shown as a transparent ribbon for clarity. **B)** Skp supports the solubility of LipA over a broad range of ionic strength. The soluble fraction of the wild-type LipA in presence of Skp (light blue) was determined from SDS-PAGE after the sedimentation assay at various salt concentrations. The assays were performed in technical triplicates, the mean values and SD are shown. Salt-dependent aggregation of LipA alone and LipA in presence of LipH are indicated in red and dark blue, respectively, and correspond to those in Figure 2B. **C)** Microscale thermophoresis of LipA_F144E_:Skp interaction, using fluorescently labelled LipA_F144E_ and Skp titrations (monomers concentration indicated). The assay was performed in technical triplicates, the mean values and SD are shown. **D)** SEC-MALS of the LipA_F144E_:Skp_3_ complex and Skp_3_ alone. Normalized UV absorbance of LipA_F144E_:Skp_3_ (solid black line) and Skp_3_ (dotted black line), and the corresponding molecular weights are indicated. Below: SDS-PAGE of the LipA_F144E_:Skp_3_ eluate. **E)** Solubility of the lipase is dependent of the LipA:Skp_3_ ratio. The soluble fraction of the wildtype LipA was determined from SDS-PAGE after the sedimentation assay at various LipA:Skp_3_ ratios. The assays were performed in technical triplicates, the mean values and SD are shown.

Next, we set out to determine the binding affinity of Skp to LipA. Earlier studies of Skp interactions with its most abundant clients, OMPs, provided the affinity estimates at low-nanomolar to low-micromolar scales for OMP:Skp_3_ complexes formed at 1:1 and 1:2 stoichiometry, respectively ^45–48^, while the interactions with soluble proteins have not been addressed. For the analysis here, we employed microscale thermophoresis (MST) and used the aggregation-resistant mutant LipA_F144E_ as the fluorescently labelled component. LipA_F144E_-FM was mixed with Skp ranging from 4.5 nM to 37 μM (concentration of Skp monomers), and the MST response of LipA_F144E_-FM, i.e. its mobility within the micrometer-sized temperature gradient, was analysed. Strikingly, LipA_F144E_ manifested a bimodal MST response upon increasing the Skp concentration that appeared as a V-shaped plot with a transition at approx. 300 nM Skp_1_ (100 nM Skp_3_) (Figure 7C). The data suggested two distinct modes of binding to the chaperone, where individual K_D_’s were ~20 nM and 2 μM, and the values were in a good agreement with values measured for OMP:Skp_3_ interactions where either one or two Skp timers were involved ^45–47^.

The LipA:Skp affinity measured in the MST experiments suggested that two copies of Skp trimer were recruited to stabilize the soluble lipase (Figures 4 and 5; monomeric Skp of 15 μM). To test the stoichiometry of the complex experimentally, SEC-MALS analysis was performed on LipA_F144E_:Skp mixture (molar ratio 1:3; Figure 7D). The elution peak demonstrated a prominent shift in comparison to Skp_3_ alone, and the estimated molecular mass of 139 kDa approached closely the calculated mass of the complex LipA:2*Skp_3_ (148 kDa). SDS-PAGE confirmed presence of both the chaperone and the lipase in the peak fractions, and the densitometry analysis of the corresponding bands suggested the excess of Skp (estimated LipA:Skp_3_ ratio 1:3). The remaining free LipA fraction was likely bound to the column resin, as we observed previously. Thus, we concluded that the relatively small protein LipA provided sufficient surface area for binding two copies of the trimeric Skp. To test, whether this stoichiometry was beneficial for the solubility of the lipase, we used the aggregation-prone wild-type LipA in the sedimentation assay where the LipA:Skp_3_ ratio was varied. Strikingly, the solubility of LipA dropped dramatically when LipA:Skp_3_ molar ratio was reduced to 1:1 (monomeric Skp 1.5 μM), suggesting that a single Skp trimer was not capable of rescuing the client from aggregation/misfolding. It seems plausible that at elevated Skp concentrations, two copies of the chaperone lock LipA within the consolidated cavity, and rapid re-binding of the escaped clients occurs prior to the irreversible aggregation.

## Discussion

The protein fold is determined by its primary amino acid sequence, but the pathways followed by the polypeptide chain towards the functional structure commonly involve the assistance of chemical and proteinaceous chaperones. The chaperones steer their client dynamics along the energy landscape, facilitate the local and global folding/unfolding events, prevent off-pathway intermediates, and serve to disaggregate occasional misfolded structures ^49,50^. The role of chaperones is of particular importance under stress factors, such as elevated temperature, or in challenging environments, such as bacterial periplasm, where the protein folding should proceed under broadly varying conditions. Here, we demonstrate that the nonspecific periplasmic chaperone Skp of the pathogenic bacterium *P. aeruginosa* efficiently protects the secretory lipase LipA from aggregation and suppresses its assembly into amyloid-type structures. The structural analysis of trimeric Skp in solution by SAXS reveals an ensemble of co-existing states. The interaction analysis of the LipA:Skp_3_ complex suggests that either one and two copies of Skp trimers may bind the lipase via hydrophobic interactions, but two copies are required for stabilizing the aggregation-prone protein.

Folding of the lipase LipA takes place in the periplasm, where the protein is transiently localized and interacts with the specialized chaperone LipH. The chaperone serves for correct positioning of the lid domain of LipA, thus rendering the active enzyme prior its further secretion via T2SS ^14,15^. The non-activated lipase tends to acquire loosely folded intermediate states, which are accessible for proteases and prone to aggregation ^18,51^. The chaperone-dependent folding is not common among bacterial lipases, and it might be related to the particular structure of the lid domain ^52^. In agreement with the earlier findings of Chiti and co-workers ^29,51^, our data show that the lid domain greatly contributes to LipA aggregation and misfolding into β-stranded structures, thus interfering with the protein biogenesis. As the assembly of the LipA:LipH complex rescues LipA from aggregation to a large extent, two distinct functions can be assigned to LipH, namely stabilization and activation of the client lipase. Speculatively, the chaperone might initially serve to stabilize the lipase in its folded state and hand it over to T2SS, and through the course of the evolution it has gained the propensity to activate the lipase via steric re-positioning of the lid domain.

Several challenges can be envisioned for productive LipA:LipH interactions in the living cell. Firstly, the expression levels of the foldase are substantially lower than those of LipA, so each LipH molecule should carry out multiple chaperoning cycles upon binding nascent lipases, refining the structure and releasing them for further secretion ^14^. Secondly, the accessibility of the membrane-anchored foldase for its client may be limited, especially in the crowded periplasmic environment. Finally, the elevated ionic strength encountered in the sputum of cystic fibrosis patients^31,53^ strongly promotes the aggregation of LipA and may also inhibit the electrostatic interactions at the LipA:LipH interface. Thus, accessory factors, such as non-specific but ubiquitous periplasmic chaperones may be beneficial for biogenesis of the intrinsically unstable lipase. Here, we elucidate that Skp_3_, a conditionally essential protein in *P. aeruginosa* ^54^, is a potent holdase for LipA that efficiently maintains the soluble state of the lipase and blocks the assembly of β-structured aggregates. Notably, even 10-fold excess of Skp_3_ did not inhibit LipA activation and thus allowed LipA:LipH interactions, so we described the chaperone as a putative factor in LipA biogenesis that steers folding pathways by preventing aggregation events.

Up to date, the vast majority of insights on Skp functioning have been gained from studies on the chaperone from *E. coli*. Differently from other periplasmic chaperones, Skp targets its clients based on the hydrophobicity exposed in the unfolded/misfolded structures rather than a specific sequence motif or conformations of proline residues ^46,55^. The periplasmic concentration of *Ec*Skp may reach hundreds of μM ^9^, and the chaperone extensively interacts with unfolded OMPs ^46,47,56^. Although no OMP essentially depends on Skp in the presence of SurA ^56,57^, other moonlighting activities have been suggested exclusively for Skp. Recently, the chaperone has been described as an adaptor protein that delivers the misfolded OMP LamB to the periplasmic protease DegP ^58^. Skp is a potent disaggregase of another OMP, OmpC ^59^ and it also demonstrates the holdase and disaggregase activity towards several soluble proteins ^25,26,60^. Most notably, heterologous expression and secretion of a lipase from *Burkholderia* could be substantially improved in *E. coli* upon co-expression of Skp, but not SurA ^27^, and the Skp-mediated solubility of the lipase was proposed as the decisive factor, in agreement with our direct biochemical evidences. While the functional insights on Skp of *P. aeruginosa* are sparse ^61^, the chaperone appears to be essential for the survival of bacteria in the sputum media, where the salt concentration exceeds 100 mM ^54^.

Structural analysis on Skp_3_ of *P. aeruginosa* suggests that its architecture and dynamics resemble those of *Ec*Skp_3_, despite the low sequence similarity and presence of multiple proline residues. *P. aeruginosa* Skp exists as an assembled trimer in solution, and its conformational plasticity and the hydrophobic interior fulfil the requirements for being a promiscuous nonspecific holdase chaperone. The SAXS data describe the transient outward motion of the “arm” domains that ensures opening of the interior cavity of the Skp trimer. The aggregation-prone folding intermediates of LipA are likely to be recognized by Skp_3_, kept in the protective hydrophobic cavity and then handed over to LipH for activation.

Using the aggregation-resistant lipase mutant LipA_F144E_ allowed estimating its affinity to Skp. MST measurements revealed a complex biphasic binding with affinities of ~20 nM and 2 μM. These affinities are in a remarkable agreement with the values measured for Skp interaction with various OMPs, where binding of either one or two copies of Skp_3_ has been described ^45–48^. The close match between the Skp affinities to different classes of clients is likely due to non-specific hydrophobicity-based interactions. Unexpectedly though, the relatively small protein LipA is able to recruit two trimers of Skp, suggesting that the exposed hydrophobic areas are sufficiently large to form the extensive binding interface.

Our data provide first evidence for selective interactions of *P. aeruginosa* LipA with the general and highly abundant periplasmic chaperone Skp that result in stabilization of the lipase. It remains to be shown to what extent Skp is involved in LipA biogenesis in *P. aeruginosa*, and whether the chaperone:client interactions are beneficial at particular conditions, such as elevated ionic strengths. Future experiments in the native environment of *P. aeruginosa* combined with *in vitro* analysis will account for the membrane-anchored LipH and its putative competition with Skp for the nascent LipA, as well as the handover of LipA to the T2SS machinery and potential involvement of Skp at that stage.

## Materials and methods

### Molecular cloning

The genes of *P. aeruginosa* PAO1 strain were identified using Pseudomonas Genome Database (www.pseudomonas.com). Those included PA2862 (*lipA*), PA2863 (*lipH*), PA3262 (*fkpA*), PA3801 (*yfgM*), PA1805 (*ppiD*), PA0594 (*surA*) and PA3647 (*skp/ompH/hlpA*). The genomic DNA from *P. aeruginosa* PAO1 served as a template for the amplification of the genes of interest via PCR using the KOD Xtreme polymerase (Novagen/Merck) and cloning primers containing the restriction sites (Eurofins Genomics). The PCR products were isolated using the NucleoSpin Gel and PCR Clean-up kit (Macherey-Nagel). Standard molecular cloning techniques were further employed to insert the genes of interest into target vectors using restriction nucleases (New England Biolabs). The amplified genes encoding the full-length periplasmic chaperones were initially cloned into the pET21a vector to contain C-terminal hexa-histidine tags. The following combinations of the restriction sites were used for cloning: *BamHI/HindIII* for *skp and surA, EcoRI/HindIII* for *yfgM* and *ppiD*, and *EcoRI/SalI* for *fkpA*. The positions of signal peptides and membrane-anchoring domains within the chaperones were identified using SignalP-5.0 and TMHMM services, respectively ^62,63^. For overexpression of the soluble proteins, the regions encoding the N-terminal signal peptides and the membrane anchors were removed via PCR and the plasmids were re-ligated. *E. coli* strain DH5α (Thermo Fisher Scientific) served as a recipient strain for cloning and plasmid multiplication. Plasmid DNA was isolated using the NucleoSpin Plasmid kit (Macherey-Nagel) and analysed by Sanger sequencing (Eurofins Genomics).

### Expression and purification of periplasmic chaperones

The soluble chaperones were heterologously expressed in *E. coli* BL21(DE3), and Skp was expressed in *E. coli* BL21(DE3)pLysS. Overnight cultures were grown at 37 °C upon shaking at 180 rpm and used for inoculation of pre-warmed LB medium. Cells were grown at 37 °C upon shaking at 180 rpm until reaching OD_600_ 0.6, and overexpression was induced with 0.5 mM isopropyl-β-D-thiogalactopyranoside (IPTG; Merck/Sigma-Aldrich). After 2 hours expression the cells were harvested by centrifugation at 6000 g for 15 min, 4°C (rotor SLC-6000, Sorvall/Thermo Fischer) and resuspended in 50 mM KCl, 1 mM DTT, 10 % glycerol, 20 mM HEPES pH 7.4 supplemented with protease inhibitors (cOmplete Protease Inhibitor Cocktail, Roche). The cells were lysed by shear force (M-110P cell disruptor, Microfluidics Inc.) and the debris and membranes were removed by ultracentrifugation at 205,000 g for 1 hr at 4 °C (rotor 45 Ti, Beckman Coulter). The supernatants were applied to IMAC with Ni^2+^-NTA-agarose resin (Qiagen) in the presence of 5 mM imidazole. After binding and extensive wash with the resuspension buffer containing 10 mM imidazole to remove weakly and non-specifically bound proteins, the chaperones were eluted with the buffer supplemented with 300 mM imidazole. The elution fractions were concentrated and subjected to SEC using Superdex 200 Increase 10/300 GL column (Cytiva). Fractions containing the chaperones were identified by SDS-PAGE, and the protein concentrations were determined spectrophotometrically based on calculated extinction coefficients at 280 nm: SurA 30940 M^−1^ cm^−1^, FkpA 8940 M^−1^ cm^−1^, PpiD 24870 M^−1^ cm^−1^, YfgM 8940 M^−1^ cm^−1^ and Skp 5960 M^−1^ cm^−1^. Monomer protein concentrations were adjusted to 50 μM and the aliquoted proteins were flash frozen and stored at −80°C. Before experiments, the samples were thawed, centrifuged at 100.000 g, 4°C for 1 h to remove occasional aggregates and applied for SEC to match buffering conditions required for the conducted assays.

The lipase-specific foldase LipH lacking the TMD and bearing the N-terminal hexa-histidine tag was expressed heterologously as a soluble protein in *E. coli* BL21(DE3) using plasmid pEHTHis19 ^64^. LipH was purified by IMAC and subsequent SEC in 100 mM NaCl, 200 μM TCEP, 10% glycerol and 50 mM Tris-HCl pH 8.0. Samples were flash-frozen and stored at −80 °C. Directly before experiments LipH samples were thawed, centrifuged at 100.000 g, 4 °C for 1 h and subject for SEC in 100 mM NaCl, 10 % glycerol and 20 mM Tris-HCl pH 8.0 unless other buffers are specified.

### Size exclusion chromatography combined with multi-angle light scattering

SEC-MALS was employed to probe the oligomeric states of the isolated chaperones of P. aeruginosa. The purified proteins were concentrated to 2 mg/mL using centrifugal filters with either 3 kDa or 30 kDa cut-off (Amicon Ultra-0.5, Merck/Millipore) and the samples were centrifuged at 100,000 g, 4 °C for 1 h to remove occasional aggregates. Subsequently, 80-200 μL were injected on Superdex 200 Increase 10/300 GL column (Cytiva) connected to miniDAWN TREOS II light scattering device and Optilab-TrEX Ri-detector (Wyatt Technology Corp.). FkpA, SurA and PpiD were analysed in 50 mM KCl, 5 mM MgCl_2_ and 20 mM HEPES-KOH pH 7.4; YfgM in 5 mM glycine, 20 mM NaCl and 5 mM Tris-HCl pH 8.5; Skp in 100 mM NaCl and 20 mM Tris-HCl pH 8.0, and LipH in 100 mM NaCl, 200 μM TCEP and 50 mM Tris-HCl pH 8.0. The data analysis was performed with ASTRA 7.3.2 software (Wyatt Technology Corp.).

### Small-angle X-ray scattering

SAXS data on Skp chaperone was collected using Xeuss 2.0 Q-Xoom system from Xenocs, equipped with a PILATUS3 R 300K detector (Dectris) and a GENIX 3D CU Ultra Low Divergence X-ray beam delivery system. The chosen sample-to-detector distance for the experiment was 0.55 m, resulting in an achievable q-range of 0.05 - 6 nm^−1^. The measurement was carried out at 15° C with a protein concentration of 1.20 mg/mL. Skp samples were injected in the Low Noise Flow Cell (Xenocs) via autosampler. 24 frames were collected with an exposure time of ten minutes per frame and the data was scaled to the absolute intensity against water.

All used programs for data processing were part of the ATSAS software package, version 3.0.3 ^65^. Primary data reduction was performed with the program PRIMUS ^66^. With the Guinier approximation, the forward scattering *I(0*) and the radius of gyration (*R*_g_) were determined. The program GNOM ^67^ was used to estimate the maximum particle dimension (**D*_max_*) with the pairdistribution function *p(r*). The *ab initio* reconstitution of the protein structure by dummy residues with P3 symmetry was performed using the GASBOR program ^39^. The Skp trimer model was built using the cloud-based AlphaFold 2 Multimer algorithm ^40,41^. The conformation analysis was performed using Ensemble Optimisation Method (EOM) using the default parameters (10000 models in the initial ensemble, native-like models, constant subtraction allowed) ^43,44^.

### Lipase isolation

The gene PA2862 encoding for the mature LipA was cloned into pET22b plasmid via *NdeI/BamHI* restriction sites ^64^ and the protein was expressed in *E. coli* BL21(DE3). Briefly, the overnight culture with LB ampicillin (100 μg/mL) grown at 37°C upon shaking at 180 rpm used to inoculate 100 mL LB media. Cells were grown at 37°C upon shaking at 180 rpm till reaching OD_600_ 0.6. LipA expression was induced by addition of 0.5 mM IPTG to the culture and was conducted for 2 h at 37 °C. Cells were harvested by centrifugation at 4000 g, 4°C for 15 min and resuspended in 50 mM Tris-HCl pH 7.0, 5 mM EDTA, 1 mM TCEP, supplemented with 10 μg/mL DNAse I and 50 μg/mL lysozyme (buffer IB). The suspension was incubated at 20 °C for 15 min and vortexed briefly, and the cells were lysed via sonication (ultrasonic homogenizer UP100H equipped with MS7 tip). The inclusion bodies were sedimented by centrifugation at 15000 g, 4°C for 10 min and suspended in the buffer IB for washing, followed by centrifugation. After repeating the procedure twice, the pellet was washed with 50 mM Tris-HCl pH 7.0. The inclusion bodies were dissolved in 8 M urea and 20 mM Tris-HCl pH 7.25, and the insoluble material was removed via a centrifugation step (21000 g, 10 min, 4°C). The protein purity was assessed by SDS-PAGE and subsequent staining (Quick Coomassie stain, Serva). LipA concentration was determined spectrophotometrically (extinction coefficient at 280 nm, ε_280_ = 27515 M^−1^ cm^−1^). LipA was aliquoted in reaction tubes (Low Protein Binding, Sarstedt), flash-frozen and stored at −80°C.

Optionally, LipA was fluorescently labelled to increase the detection efficiency in the sedimentation assay. LipA contains two endogenous cysteines in positions 183 and 235, which were targeted for site-specific fluorescent labelling. The urea-denatured LipA was incubated in 25-fold molar excess of fluorescein-5-maleimide (FM, Thermo Fisher Scientific) for 3 h at the ambient temperature. After the incubation, LipA was precipitated with 15 % TCA for at least 1 h on ice. The precipitated proteins were sedimented via centrifugation at 21000 g, 4°C for 15 min, and the supernatant was removed. The pellets were washed with 0.5 mL ice-cold acetone, and then repeatedly sedimented via centrifugation at 21000 g, 4°C for 10 min. After drying at 37 °C, LipA-FM pellet was solubilised in 8 M urea and 20 mM Tris-HCl pH 7.25. To remove the remaining free dye, TCA precipitation and washing steps were repeated twice. The labelling efficiency of ~150 % was determined spectrophotometrically based on the absorbance at 280 and 495 nm and the molar extinction coefficients for LipA and FM, respectively. LipA-FM was visualised on SDS-PAGE via blue-light excitation and following Coomassie blue staining.

### *In vitro* activity of the lipase

To measure the hydrolytic activity of LipA, equimolar concentrations of the foldase LipH in TGCG buffer (5 mM Tris pH 9, 5 mM glycine, 1 mM CaCl_2_, 5% glycerol) and the urea-denatured LipA (8 M urea and 20 mM Tris-HCl pH 7.25) were mixed together at concentration 1 μM in TGCG buffer to form lipase:foldase complexes in reaction tubes (Low Protein Binding, Sarstedt). The complex formation was set for 15 min at 37°C. After the incubation, 10 μL of the sample were diluted 10-fold with the TGCG buffer in 96-well plates. For the substrate preparation, 10 mM para-nitrophenyl butyrate, ~1.76 μL, (*p*NPB, Merck/Sigma) were diluted from the stock solution in 1 mL acetonitrile. Immediately before starting the measurement, the substrate solution was diluted 10-fold with 50 mM triethanolamine pH 7.4 and mixed. Subsequently, 100 μL of the substrate solution was transferred to 96-well-plates with the preassembled LipA:LipH complex, so each well contained 0.5 mM *p*NPB and 50 nM LipA:LipH complex. For the negative control, LipA without LipH were treated the same as all other samples. To measure the autohydrolysis of the substrate, both LipA and LipH were omitted, but the corresponding buffers were added to *p*NPB. The hydrolysis of *p*NPB to *p*-nitrophenolate and butyric acid was determined by measuring the absorbance of the liberated p-nitrophenolate at 410 nm over 3.5 h at 37°C on a plate reader (Infinite 200 pro, TECAN). Samples were shaken for 5 sec prior to each measurement. The hydrolytic activity of LipA was analysed by monitoring the hydrolysis based the p-nitrophenolate absorbance (as previously described) and by calculating the active LipA in U/mL in the linear range of the reaction (first 15 min), with the estimated molar extinction coefficient of p-nitrophenolate at 410 nm under the applied conditions of 12500 M^−1^ cm^−1 68^. The light path length in the well was experimentally determined and was of 0.53-0.55 cm upon applied conditions.

When indicated, the experiments were performed in the presence of NaCl and calcium upon LipA:LipH assembly and during the substrate hydrolysis measurements. Prior to the experiments, LipH was transferred into 100 mM NaCl, 10 % glycerol and 20 mM Tris-HCl pH 8.0 by SEC, and the composition of the buffer for LipA:LipH assembly, as well as for hydrolysis measurements were adjusted. To study the effect of the periplasmic chaperones (Skp, YfgM, FkpA, SurA, PpiD) on the hydrolytic activity, the chaperones were transferred into either the TGCG buffer or 100 mM NaCl, 10 % glycerol (v/v) and 20 mM Tris-HCl pH 8.0 by SEC. The proteins, LipA and LipH, were added in equimolar concentrations (1 μM) to form the complex, while the periplasmic chaperones were added in 5-fold molar excess. The complex formation and hydrolysis measurements were performed as described above. The activation of LipA by the periplasmic chaperones was also determined by adding LipA to the periplasmic chaperones at 1:5 molar ratio in the absence of LipH. To probe the effect of the periplasmic chaperones on LipA aggregation prior the activation, 1 μM LipA was incubated with individual periplasmic chaperones for 15 min at 37°C. Afterwards, 1 μM of LipH was added to the mixtures and incubated for 15 min at 37°C. The measurements of pNBP hydrolysis were performed as described above.

### Sedimentation analysis of LipA aggregation

LipA aggregation at various conditions was probed by sedimentation assay. To improve the detection sensitivity and so facilitate the experiment at low lipase concentrations, site-specific fluorescent labelling of the lipase was introduced. The endogenous cysteines at positions 183 and 235 were conjugated with FM, as described above, so as little as 5 ng LipA-FM could be detected via in-gel fluorescence imaging. The urea-denatured LipA-FM was diluted with the TGCG buffer from 150 μM stock solution to 1.6 μM and kept on ice protein in reaction tubes (Low Protein Binding, Sarstedt). The chaperones were transferred to 1.5 mL polypropylene reaction tubes containing 20 mM Tris-HCl pH 8.0 and varying NaCl concentrations, to achieve the specific ionic strength indicated for each assay. The minimal ionic strength conditions were probed when using the TGCG buffer. 15.6 μL of LipA-FM were added to the tube and mixed by pipetting, so the reactions contained 0.5 μM LipA-FM and 5 μM of individual chaperones in the total volume of 50 μL. For the dimeric FkpA and trimeric Skp, the monomer concentrations were 10 μM and 15 μM, respectively. The LipA-specific foldase LipH was used in the equimolar ratio to the lipase (final concentration 0.5 μM). The reaction tubes were incubated at 37 °C for 15 min to promote LipA aggregation and the samples were then centrifuged (21000 g, 15 min, 4 °C) to sediment the aggregates. To avoid disturbing the pellets, 40 μL of each supernatant fraction were transferred to new 1.5 mL tubes. The remaining material contained the aggregated LipA as a pellet and also 10 μL of the supernatant fraction. Both samples were precipitated by adding 100 μL of 20 % TCA and incubating for 15 min on ice. The precipitated proteins were pelleted upon centrifugation at 21000 g, 4°C for 10 min, TCA was removed, and the samples were washed, as described above. Further, 15 μL of 2x reducing SDS-PAGE sample buffer were used to wash the tube walls, and 5 μL of the sample were loaded on SDS-PAGE and in-gel fluorescence was recorded. The signal was quantified by ImageQuant software (Cytiva). The background was subtracted using the local average algorithm. The signals of the supernatant (*I*_sol_) represent the soluble fraction of LipA and correspond to 80 % of the total soluble LipA. The value was used then to calculate the actual signal intensities of the soluble (*I*_sol, corr_) and aggregated (*I*_agg, corr_) LipA, as:

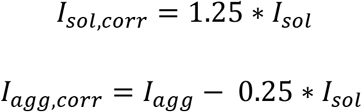

where *I*_agg_ is the signal measured for the aggregated LipA and the remaining 10 μL of the supernatant. The soluble fraction of LipA was calculated as a ratio:

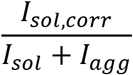

Each assay was carried out in triplicates.

### Amyloid-specific ThT fluorescence

Thioflavin T (ThT, Merck/Sigma-Aldrich) was dissolved to the concentration of 100 μM in the buffer TGCG, and the solution was kept on ice. Immediately prior to the experiment, ThT was diluted with the TGCG buffer to 10 μM and mixed with 0.5 μM urea-denatured LipA and 5 μM chaperones. For the dimeric FkpA and trimeric Skp, the monomer concentrations were 10 μM and 15 μM, respectively. The total volume was set to 65 μL and the reaction was carried out at room temperature in reaction tubes (Low Protein Binding, Sarstedt). For the control experiments, containing either the chaperones or LipA, the reaction volume was adjusted to 65 μL with an appropriate buffer. After incubation at 37 °C for 15 min, each reaction was transferred into a quartz cuvette for measuring ThT fluorescence at the fluorescence spectrophotometer (Fluorolog 3, Horiba Scientific). The excitation wavelength was set at 450 nm, and the emission spectra were recorded from 467 to 520 nm. ThT fluorescence intensity at 485 nm was used to evaluate and compare LipA aggregation between samples. Each measurement was carried out in independent triplicates.

### Microscale thermophoresis

MST was used to monitor interactions of Skp from *P. aeruginosa* with FM-labelled LipA_F144E_. LipA_F144E_ was diluted to 100 nM in the TGCG buffer supplemented with 0.05% Tween 20 and kept on ice protected from light. For the MST measurement, 10 μl of 50 nM LipA were mixed with Skp ranging from to 2.3 nM to 18.5 μM in a 0.5 mL reaction tube (Low Protein Binding, Sarstedt). LipA:Skp samples were incubated for 15 min at 22 °C in the dark, then loaded into Premium-type capillaries and analysed in Monolith NT.115 instrument (NanoTemper Technologies, Munich, Germany). The MST power was set to 80 % and the LED power in the blue channel was set to 60 %. The thermophoresis was detected by the normalised fluorescence time traces for 30 seconds with 5 seconds delay and 5 seconds for recovery. The putative LipA aggregation at the capillary surface was controlled by recording the fluorescence intensity profiles of individual capillaries before and after the experiment (time difference approx. 1 hour). The data was evaluated by NT Analysis software (NanoTemper Technologies, Munich, Germany). Each sample was analysed twice, and the measurement were performed in independent triplicates.

### Transmission electron microscopy

TEM was utilized to visualize the formation of LipA aggregates. All buffers were filtered with 0.1 μm filters (Whatman Puradisc 0.1 micron). Denatured LipA was diluted to 300-500 μM in 20 mM Tris-HCl pH 7.25 and 8 M urea. In case of TEM analysis of LipA in urea, the protein was further diluted in 20 mM Tris-HCl pH 7.25 and 8 M urea. For urea-free measurements, LipA was diluted to 3-20 μM in either TGCG buffer or in 150 mM NaCl, 10 % glycerol and 20 mM Tris-HCl pH 8.0. Samples containing Skp or LipH were prepared in 20 mM Tris-HCl pH 8.0, 150 mM NaCl and 10 % glycerol to promote lipase aggregation and visualize the anti-aggregating effect of the chaperones. LipA:LipH samples contained 5 μM LipA and 15 μM LipH (molar ratio 1:3) and LipA:Skp_3_ samples contained 3 μM LipA and 15 μM Skp_3_ (molar ratio 1:5). All samples were incubated for 15 min at 37°C and used for grid preparation on a carbon-coated copper grid. 3 μL of sample were added on top of the grid incubated for 1 min at the ambient temperature. Excess liquid was removed using filter paper. For negative staining, the grid was incubated in 2 % uranyl acetate (solution in distilled water pH 4.3) for 1 min in the dark. Afterwards, excess liquid was removed. Grids were dried at the ambient temperature for at least 15 min prior TEM imaging imaging. TEM images were acquired at the Zeiss EM902 operating at 80 KV using a slow-scan CCD-Camera (Typ 7899 inside) controlled by the imaging software ImageSP (SYSPROG, TRS) and prepared using the image processing software “Fiji” ^69^.

## Data availability

We uploaded the SAXS data to the Small Angle Scattering Biological Data Bank (SASBDB; www.sasbdb.org) with the accession code SASDNM5.

## Contributions

AP and MBu carried out cloning, protein expression and isolation, and biochemical and fluorescence-based experiments; JR and SHJS carried out SAXS measurements and structural characterization of chaperones; EH carried out SEC-MALS analysis; MBä carried out TEM imaging; LS, KEJ, FK, SHJS and AK supervised the research. AP and AK wrote the manuscript with help of all co-authors.

## Acknowledgements

The research was supported by the German Research Foundation (Deutsche Forschungsgemeinschaft, DFG) via the Collaborative Research Centre 1208 “Identity and Dynamics of Membrane Systems - from Molecules to Cellular Functions” (project 267205415, sub-projects A1, A2, A10, and Z02). The Center for Structural Studies is funded by DFG, projects 417919780 and INST 208/761-1 FUGG. Center of Advanced imaging is funded by DFG, project 284074525. Our special thanks go to Jessica Hausmann and Stefanie Weidtkamp-Peters for their kind support on TEM.

## Competing interests

The authors declare no competing interests.

## Notes

### Competing Interest Statement

The authors have declared no competing interest.

